# PacBio genome assembly of *Olea europaea* L. subsp. *europaea* cultivars ‘Frantoio’ and ‘Leccino’ reveal main structural differences in key genes related to salt stress

**DOI:** 10.1101/2024.12.13.628337

**Authors:** Luca Sebastiani, Iqra Sarfraz, Alessandra Francini, Mirko Celii, Rod A. Wing, Andrea Zuccolo

**Affiliations:** Institute of Crop Science, Scuola Superiore Sant’Anna, Piazza Martiri della Libertà 33, 56127, Pisa; King Abdullah University of Science & Technology (KAUST), Saudi Arabia Andrea Zuccolo, Mirko Celii, Rod A. Wing; Arizona Genomics Institute, School of Plant Sciences, University of Arizona, Tucson, AZ, USA Rod A. Wing

## Abstract

We present two high-quality genome assemblies for *Olea europaea* L. cultivars ‘Frantoio’ and ‘Leccino’ leveraging PacBio HiFi sequencing to achieve approximately 30× genome coverage for each cultivar. The assemblies span 1.18 Gbp and 1.43 Gbp with contig N50 values of 1.78 Mbp and 45.88 Mbp for ‘Frantoio’ and ‘Leccino’ respectively. BUSCO analysis revealed a great genome completeness (∼97.9%), surpassing many of earlier *Olea europaea* assemblies and is in par with the most recent one. Repetitive content accounted for ∼67.5% in ‘Frantoio’ and ∼70.8% in ‘Leccino’ with long terminal repeats (LTRs) dominating. Notably, a tandem repeat family, Satellite 1, represented ∼16.9% and ∼8.6% of the ‘Leccino’ and ‘Frantoio’ genomes, respectively. The structural variant (SV) analysis was done with a particular focus on those associated with nine key gene families involved in salinity tolerance and identified cultivar-specific genomic differences, emphasizing the diversity within domesticated olives. This comprehensive analysis provides valuable resources for studying olive genome evolution, domestication, and genetic improvement, underscoring the utility of long-read sequencing for resolving complex genomic features.

## Background & Summary

Olive (*Olea europaea* L. subsp. *europaea*) is an economically relevant and widely distributed fruit crop in the Mediterranean Basin. This iconic tree has its origins linked to the beginning of ancient Mediterranean civilizations dated more than six millennia ago (**1**) and its domestication is still debated in scientific literature (**2**). Thanks to the healthy and organoleptic properties of the extra virgin olive oil, in the last decades olive orchards spread in many other warm-temperate regions of the world such as North and South America, Australia, New Zealand, and South Africa, and even in the monsoon areas likes China and India.

Despite the environmental, cultural, economic and scientific value of this specie, the availability of a high-quality genome data for relevant cultivars are still scarce. Additionally, the selection of olive varieties using traditional breeding practices is a time consuming and largely random process, since precise molecular information on genes location and structure are largely missing. Today, third generation sequencing techniques (**3**) generating very long reads allow the high-quality complete assembly of complex genomes. These genome assemblies in turn enable a deeper understanding of genome structure and provide foundational data sets for functional genomics, genetic engineering and molecular breeding. All these aspects are particularly relevant in woody crops, like olive, where genome data are missing, scarce, or when present are often characterised by lower quality in comparison to other plants/crops. Additionally, olive has a complex mid-size genome characterized by high heterozygosity and high repeat content.

In olive, the first research aimed to achieve deeper genomic information was done using Sanger and 454 pyrosequencing technologies and targeted transcriptome (**4**). Using this approach, 2 million reads from 12 cDNA libraries were attained from several olive cultivars and progenies. These libraries were done for fruit in different developmental stages, vegetative organs (stems, leaves, roots) and buds. In 2016 (**5**) using a blend of fosmid and whole genome shotgun libraries and Illumina sequencing technology, sequenced the genome of a single 1200-year-old Mediterranean olive tree (*Olea europaea* L. subsp. *europaea* var. *europaea* cultivar ‘Farga’). This genome assembly has a total length of 1.31 Gb that correspond to 95 % of the estimated 1.38 Gb Olive genome size and about 56,349 unique protein coding genes were predicted. This first assembled draft genome of *Olea europaea* was a milestone for the study of the evolution and domestication processes of olive and gave new insight in the genetic bases of key phenotypic traits relevant to agronomic and stress tolerance characteristics (**6**). More recently the olive cultivar Arbequina, suitable for mechanized harvesting and dense planting, was sequenced using Oxford Nanopore third generation sequencing (**7**). Authors assembled 1.1 Gb of sequences organized in 23 pseudochromosomes and predicted 53,518 protein-coding genes. The greater contiguity of this genome assembly allowed the identification of 202 genes part of the oleuropein biosynthesis pathway genes which is twice the number of those identified from previous genomic data. More recently, (**8**) provided a gapless genome assembly for *O. europaea* cultivar ‘Leccino’ exploiting that resource and transcriptomic and metabolomic data to unvail a pivotal regulatory mechanism in oil biosynthesis. This evidence once more underlines the advantages provided by high-quality genome assembly for genomics studies. The accessibility of high-quality genome sequences today provides an unprecedented opportunity to compare cultivars, enabling the identification of genetic variations underlying traits such as yield, disease resistance, and environmental adaptation. Building on this potential, we targeted the high-quality genome assembly of two important olive cultivars, ‘Leccino’ and ‘Frantoio’, to gain deeper insights into salt-stress tolerance.

Salt stress-together with droughts - is a major issue in the Mediterranean region. Furthermore, in the last decades, salinity problems are becoming more extensive due to incorrect irrigation practices, use of marginal land and negative effects of climate changes (**9**). Additionally, agricultural water use is in competition with the civil and industrial sectors lead more often to the use of poor-quality water that could contain ions, such as sodium (Na^+^) and chlorine (Cl^-^), that increase the risks of soil salinization. Soil salinity has a significant economic impact, particularly in terms of reduced yield. Olive is considered a moderately resistant species to salinity (**10**), with different cultivars exhibiting varying degrees of tolerance to salinity stress. (**11**). In this respect, phenotypic difference in salinity tolerance is well known and studied in the two worldwide cultivated Italian cultivar ‘Frantoio’ (tolerant) and ‘Leccino’ (sensitive) (**12**).

More in general, plants defence mechanisms to salinity rely to Na^+^ accumulation in roots, avoiding its translocation into the aerial parts, and to cell exclusion mechanisms, such as Na^+^ compartmentation in vacuole or cell wall (**13**). Also in olive, tolerance traits have been associated with the ability to retain Na^+^ and Cl^-^ in the root, limiting xylematic translocation and transport in the aerial organs (**10**). Literature data showed that more tolerant is the cultivar less Na^+^ is translocated into the aerial part (**14; 11**) with a decreasing gradient of Na^+^ and Cl^-^ from the base to the apex of the shoot. In experimental studies using 120 mM NaCl, ‘Leccino’ (sensitive) showed a significant lower shoot Na^+^ exclusion than ‘Frantoio’ (tolerant) (**10**).

In recent years, attempts to clarify the genetic base of salinity tolerance in olive have been made studying several genes that have a role in salt tolerance (**15**). Authors found that the Na+/H+ antiport pumps of the tonoplast (*NHX*) and vacuolar ATPase subunit E (*VHA sub. E*) genes were significantly overexpressed both in ‘Frantoio’ and ‘Leccino’ treated with 120 mM NaCl. On the contrary, plasma membrane Na+/H+ antiporter (*SOS1*), *ATPase11* and *ATPase8* genes were overexpressed only in ‘Frantoio’. Further studies (**16**) showed that in ‘Leccino’ the H+-pumping expression was decreased by salinity treatment in the early phase of salt response. Overall, these data proved that genes differential activities are determinant for salinity and more relevant insight could be achieved when high quality genomic data will be available.

This study presents the genome assemblies of the olive cultivars ‘Frantoio’ and ‘Leccino’, obtained using HiFi PacBio long reads with 27X and 29X coverage, respectively, along with their baseline gene and repeat annotation, including transposable elements. Additionally, we compare these assemblies, focusing on structural variants linked to key salinity tolerance genes.

## Methods

### Sampling and genome sequencing

Young leaves were sampled from 1 year old *Olea europaea* L. cultivar ‘Frantoio’ and ‘Leccino’ plants (supplied from Società Pesciatina d’Olivicoltura and certified according to the Community Agricultural Conformity), preserved in liquid nitrogen and stored at -80 °C until subsequent analysis. High Molecular Weight (HMW) DNA was extracted using a modified CTAB protocol (**17**). The quality of the extracted DNA was assessed through pulsed-field gel electrophoresis (CHEF) on 1% agarose gels to evaluate fragment size and restriction enzyme digestibility. Quantification was performed using Qubit fluorometry (Thermo Fisher Scientific, Waltham, MA). Sequencing was performed using PacBio Revio System at the Arizona Genomics Institute (Tucson, AZ), on Revio SMRT cells (**18**). High-quality sequencing data generated, 37.5 Gbp of Hifi reads for ‘Frantoio’ and 38.7 Gbp for ‘Leccino’, with read N50 values of 5.9 kbp and 16.3 kbp, respectively. This data provided approximately 30x genome coverage for both cultivars, supporting robust genome assemblies.

### Genome size and heterozygosity estimation

The K-mer analysis was performed using Jellyfish v2.3.0 (**19**) setting the K-mer length at 31. The resulting K-mer frequency distributions were processed using Genomescope2 (**20**). The K-mer frequency distribution showed two distinct peaks (Supplementary Figure 1), the first peaks at approximately 12x-13.3x coverage, corresponded to heterozygous regions, while the second peak, at around 24x-26x coverage, reflected homozygous regions. In Frantoio, the heterozygosity rate (ab) was estimated at 2.2%, with a duplication rate (dup) of 0.0914; in ‘Leccino’ these values were 1.85% and 0.153 respectively. These heterozygosity values are higher than those reported for Arbequina i.e., 1.09% (**7**), suggesting greater diversity in ‘Frantoio’ and ‘Leccino’. The K-mer analysis indicated high repetitive content, with 51.6% and 52.2% of unique sequences ‘Frantoio’ and ‘Leccino’, respectively. The significant repetitive content in both cultivars’ genome, emphasize the need for PacBio Hifi sequencing for resolving complex regions, to achieve contiguous, accurate assemblies.

### Genome assembly and scaffolding

PacBio Hifi reads were assembled using hifiasm/v0.19.8 (**21**) with default parameters and redundant haplotigs were removed using “purge haplotigs” (**22**). The contigs contained in the primary assembly were scaffolded using the tool RagTag/v2.1.0 as described by Alonge *et al*. (**23**). For the whole-genome alignment, the built-in Minimap2 inbuilt aligner was used (**24**). The genome assembly for ‘Frantoio’ was around 1,18 Gbp with 1,726 contigs and an N50 of 1.78 Mbp, whereas ‘Leccino’ was around 1,43 Gbp with 103 contigs and an N50 of 45.86 Mbp (Table 1).

**Table 1.** Main statistics of draft genome assembly for *Olea europaea* cultivars ‘Frantoio’ and ‘Leccino’.

| Parameter | Frantoio | Leccino |
| --- | --- | --- |
| Total number of contigs ($\geq 50,000$ bp) | 1,726 | 103 |
| Total length ($\geq 50,000$ bp) | 1,184,667,663 | 1,431,741,805 |
| Largest Contigs | 7,517,611 | 73,351,808 |
| GC % | 35.81 | 36.87 |
| N50 | 1,779,939 | 45,875,387 |
| N90 | 242,455 | 6,013,538 |
| L50 | 189 | 13 |
| L90 | 850 | 34 |

**Table 2:** Scaffolding results of ‘Frantoio’ and ‘Leccino’ assemblies using ‘Farga’, ‘Sylvestris’ and ‘Leccino’ published as a reference.

|  | Sylvestris |  | Farga |  | Leccino_published |  | Leccino_assembly |  |
| --- | --- | --- | --- | --- | --- | --- | --- | --- |
|  | Coverage | Gaps | Coverage | Gaps | Coverage | Gaps | Coverage | Gaps |
| ‘Frantoio’ | 84.91 % | 598 | 37.91 % | 500 | 82.43% | 1,208 | 75.97 % | 1,408 |
| ‘Leccino’ | 80.34 % | 13 | 26.85 % | 12 | 93.37 % | 33 | - | - |

The genome size of our genome assemblies compared well with those provided by the K-mers analysis (∼1.29 Gbp for ‘Frantoio’ and ∼1.31 Gbp for ‘Leccino’) and align closely to those of the previous published olive genome assemblies such as *Olea europaea* L. subsp. *europaea* cv *‘*Farga’ (∼1.31 Gb**) (5**), Arbequina (∼1.3Gb**) (7**), and the recently released ‘Leccino’ (∼1.28 Gb) (**8**). These estimates confirm that the genome size of cultivated olive is smaller in comparison to that of wild oleaster (*Olea europaea* subsp. ‘*sylvestris’*), which is approximately 1.48 GB (**7**).

The scaffolding of the ‘Frantoio’ assembly and ‘Leccino’assembly was carried out in a stepwise approach using multiple reference genomes to enhance the assembly quality and accuracy. These references includes the wild olive (*Olea europaea* subsp. ‘*sylvestris’* RefSeq assembly GCF_002742605.1, O_europea_v1), the cultivated olive ‘Farga’ (“*Olea europaea* subsp. *europaea* genome assembly OLEA9,” 2020) and the ‘Leccino’ genome (“*Olea europaea* subsp. *europaea* genome assembly GCA-902713445.1) recently published (**8**). In each step, the ‘Frantoio’ and ‘Leccino’ (our sequencing) assembly was scaffolded independently with one reference genome at a time.

In the first step, *Olea europaea* subsp. ‘*sylvestris’* was used as a reference. The scaffold coverage achieved was 84.91% for ‘Frantoio’, and 80.34% for ‘Leccino’ (Supplementary Figure 2 and 3). A notable difference was observed in the number of structural gaps with 598 gaps in ‘Frantoio’, and 13 gaps in ‘Leccino’.

In the second step, the ‘Farga’ genome was used as a reference. Scaffold coverage decreased to 37.91% for ‘Frantoio’ and 26.85% for ‘Leccino’, both lower than the coverage achieved with ‘Sylvestris’ (Supplementary figure 4 and 5). However, structural gaps were reduced, with 500 gaps identified in ‘Frantoio’ and 12 gaps in ‘Leccino’.

In the final step, the ‘Frantoio’ genome assembly was scaffolded using the ‘Leccino’ assembly obtained in this study and published ‘Leccino’ (**8**). The scaffold coverage was 75.97% and 1,408 structural gaps with current ‘Leccino’ assembly and 82.49% and 1,028 structural gaps with ‘Leccino’ published genome (Supplementary Figure 6 and 7).

Notably, when the ‘Leccino’ assembly from this study was scaffolded with the published ‘Leccino’ genome, scaffold coverage reached 93.37% with only 33 gaps, indicating highest quality among all steps. It strongly indicates that the assembly generated in this study is of high quality and closely aligns with the published reference and substantial portion of genome is accurately captured.

### Benchmarking Universal Single Copy Orthologs (BUSCO)

BUSCO/v5.7.1 (**25**) with eudicotyledons_odb10 database, (**26**) which comprises 1,614 orthologous genes, was used to assess the completeness of the genome assembly, calculating the percentage of complete (C), complete and single copy (S), complete and duplicated (D), fragmented (F), and missing (M) genes (Supplementary Table 1).

Complete BUSCO gene copies were almost equals: 1,580 for ‘Frantoio’ and 1,581 for ‘Leccino’, corresponding to approximately 97.9% of the entire BUSCO gene set. These results are higher than the values reported in reference study (**5; 27**) with ‘Farga’ (92.99%), ‘Arbequina’ (92.87 %) and ‘Sylvestris’ (85.50%) but closer (99.93%) to that obtained by Lv *et al*. (**8**) in the cultivar ‘Leccino’. Single copy accounted for 83.1% (1,342 copies) in ‘Frantoio’ and 82.9% (1,338 copies) in ‘Leccino’. The number of duplicated ortholog was 14.7% (238 copies) for ‘Frantoio’ and 15.1% (243 copies) for ‘Leccino’ of the total BUSCO groups searched, respectively, which are comparable to Arbequina where the duplication rate was reported as 20.38% and in ‘Farga’ with 18.15% of duplication respectively and in good agreement with the 12.8% of duplicated genes reported in the cultivar ‘Leccino’ Lv *et al*. (**8**). In contrast, the wild relative *Olea europaea* subsp. *sylvestris* exhibited a much higher duplication of 37.98% (**7**).

The Fragmented orthologs accounted for 1.4% (22 copies) in both cultivars, while the missing copies were limited to 0.8% (12 copies) in ‘Frantoio’ and 0.6% (11 copies) in ‘Leccino’. These values are lower than those observed in other accessions; for example, 2.42% fragmentation is reported in ‘Arbequina’ and while ‘Sylvestris’ exhibited 6.69% fragmentation and 7.81% missing BUSCOs (**7**). This likely reflects the high contiguity and accuracy achieved in these assemblies, which reduces the incidence of fragmented and missing gene copies and enhance the completeness of the genome. Altogether the BUSCO statistics testify the high completeness and contiguity achieved in our genome assemblies.

### Repeats and Transposable Elements identification

The Extensive *de novo* TE annotator (EDTA)/v2.1.0 was used to generate a Transposable Element (TE) library (**28**). All the Helitron predictions were removed from the TE library because their identification is quite imprecise and prone to generate false positives (**28**). Following this, the RepeatMasker/v4.1.4 (**29**) was run with the default settings to mask the repetitive sequences on the genome assemblies.

The TE content of the *Olea europaea* L. assembly was estimated to be 67.47% in ‘Frantoio’ and 70.84% in ‘Leccino’. These values accounted for ∼769 Mb and ∼1,006 Mb in ‘Frantoio’ and ‘Leccino’ respectively (Table 3). This different amount of repetitive and TE related sequences explains most of the diverse genome size of the two cultivars. The TE and repetitive sequence fraction is almost in the same range of previously sequenced olive genomes, 66.30% for *Olea europaea* var.’Leccino’(**8**), 59% cultivated *Olea europaea* cv. ‘Picual’ (**30**), but significantly larger than *Olea europaea* subsp. *sylvestris*, with 51% of genome composed of repetitive DNA (**27**).

**Table 3.** Transposable elements classification according to Wicher et al. (32) in *Olea europaea* cultivars ‘Frantoio’ and ‘Leccino’.

| Class | Cultivar | Frantoio |  | Leccino |  |
| --- | --- | --- | --- | --- | --- |
|  |  | Bp mask. | % mask. | Bp mask. | % mask. |
| hAT (DTA) DNA_TE and MITE |  | 34,064,132 | 2.96 | 27,165,156 | 1.92 |
| CACTA (DTC) DNA_TE and MITE |  | 27,483,293 | 2.39 | 27,724,730 | 1.96 |
| Pif-Harbinger (DTH) DNA_TE and MITE |  | 21,040,797 | 1.83 | 22,235,407 | 1.57 |
| Mutator (DTM) DNA_TE and MITE |  | 93,016,860 | 8.10 | 121,730,286 | 8.61 |
| Tc1-mariner (DTT) DNA_TE and MITE |  | 3,703,275 | 0.32 | 2,451,638 | 0.17 |
| LTR-RT Ty1-Copia |  | 138,554,511 | 12.08 | 151,348,585 | 10.70 |
| LTR-RT Ty3-Gypsy |  | 195,143,475 | 17.01 | 212,229,332 | 15.01 |
| LTR_unknown |  | 57,157,745 | 4.98 | 55,241,398 | 3.91 |
|  | Satellite 1 | 102,869,087 | 8.62 | 242,035,084 | 16.90 |
|  | Satellite 2 | 55,565,954 | 4.65 | 82,789,861 | 5.78 |
|  | Satellite 3 | 41,014,775 | 4.53 | 61,743,874 | 4.31 |
| Total interspersed |  | 769,613,904 | 67.47 | 1,006,695,531 | 70.84 |

In both cultivar the largest TE class is represented by LTR-RT accounting for 34.07% and 29.62% of the genome size in ‘Frantoio’ and in ‘Leccino’, respectively. In both cultivars Ty3-gypsy superclass always outnumber Ty1-copia one. Altogether Class 2 DNA TE represent 15.6% and 14.23% of the genome assembly size in ‘Frantoio’ and ‘Leccino’, respectively. In both case the most abundant family seems to be that of Mutator like elements totalling 8.1% in Frantoio and 8.61% in ‘Leccino’ assembly. However, part of the elements assigned to the Mutator class could have been misclassified erroneously by EDTA and assigned to that TE families just on the bases of their structural feature. This was the case of three abundant satellite sequences we identified and named Satellite_1, Satellite_2 and Satellite_3 (Table 3, Supplementary Table 2). Satellite_1 is an 80 bp long minisatellite that represents approximately 8.62% and 16.90% of the genome assemblies of the ‘Frantoio’ and ‘Leccino’, respectively. This corresponds to about 102 Mbp in ‘Frantoio’ and 242 Mbp ‘Leccino’ genome assembly. The copy number of Satellite_1 is 1.28 million copies in ‘Frantoio’ and 3.03 million copies in ‘Leccino’. Satellite_2 is a tandem repeat characterized by a 141 bp long monomer which covers approximately 4.65% of the ‘Frantoio’ genome, corresponding to 55,565,954 bp, and 5.78% of the ‘Leccino’ one, corresponding to 82,789,861 bp. The amount of Satellite_2 copies is estimated to be 394,000 and 587,000 in ‘Frantoio’ and ‘Leccino’, respectively. Finally, Satellite_3 has a 107 bp long monomer. It covers 41.01 Mbp and 61.74 Mbp of ‘Frantoio’ and ‘Leccino’ genome assemblies, respectively. The estimated copy number is ∼383,000 in ‘Frantoio’ and ∼604,000 in ‘Leccino’. This evidence confirms the finding that a large portion of the olive genome is composed by few different families of tandemly arranged repeats (**31**).

### Gene prediction and functional annotation

The gene prediction was carried out using the tool Augustus (**33**) as implemented in the suite Omicsbox (**34**). The genome assemblies soft masked for repeats were used as the primary input and the gene structure devised in *Arabidopsis thaliana* was used as a model. The Functional annotation was done comparing the predicted genes to the nr division of GeneBank using Diamond blast /v2.1.8 and analyzing them with InterProScan/v5.61-93.0 (**36**).

Gene prediction found 59,777 genes (47,201 of high quality - posterior probability > 0.4) in ‘Frantoio’ and 36,263 of them were successfully functionally annotated. In ‘Leccino’, 67,103 genes (53,302 of high quality - posterior probability > 0.4) were found and 36,629 of them were functionally annotated. The discrepancy in the number of genes predicted in the two cultivars could likely reflect the greater fragmentation of the cultivar ‘Frantoio’ genome assembly. These values are comparable with those of Cruz *et al* (**5**), where the Authors found a set of 56,349 protein coding genes, with 89,982 transcripts encoding 79,910 unique protein products and those of the recently published cultivar ‘Leccino’ genome assembly in which 70,138 protein encoding genes were identified (**8**). In Arbequina (**7**) genome assembly, the predicted protein-coding genes were 53,518 and Authors successfully annotated 50,969 genes using GO, Kyoto Encyclopedia of Genes and Genomes (KEGG), EuKaryotic Orthologous Groups (KOG), TrEMBL, and Nonredundant (Nr) databases. Overall, gene predictions and annotations showed that in *Olea europaea* L subsp. *europaea* the protein-coding genes number range from 67,103 to 53,518, with the higher values in ‘Leccino’ and the lowest values in ‘Arbequina’.

### Structural Variant Analisys

The ‘Frantoio’ genome assembly was aligned to the ‘Leccino’ assembly using minimap2/v2.24 2 (**24**) under default parameters. The resulting sam file was sorted and indexed using samtools/v1.16.1 (**37**), subsequently, the SVIM-asm tool was used to detect SVs, including insertion, inversion, deletion, duplication, interspersed duplication and tandem duplication (**38**).

The total number of deletions, insertions and inversions in *Olea europaea* genome of cultivar ‘Leccino’ in comparison to cultivar ‘Frantoio’ are presented in Figure 1. Deletions and insertions are very similar in number (22,469 deletions and 21,218 insertions), while the number of inversions is very low (n = 33). More than 85% of the insertions and deletions exhibit high similarity to TEs or other repetitive sequences. The enrichment of TEs in structural variants, compared to the entire genome, highlights the contribution of these elements to genome variation. According to data the two cultivars ‘Frantoio’ and ‘Leccino’ exhibit a considerable amount of genetic variation, consistent with previous findings obtained using Simple Sequence Repeats (SSRs) markers (**39**).

**Figure 1.**
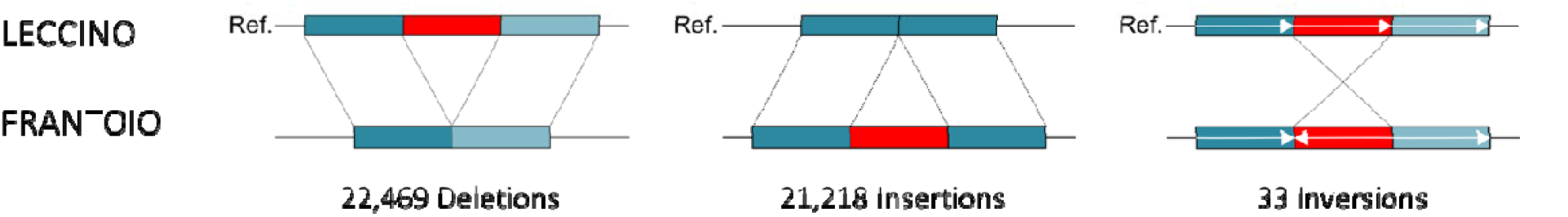
Number of deletions, insertions and inversions in *Olea europaea* genome of cultivar ‘Leccino’ (reference = Ref.) in comparison to cultivar ‘Frantoio’.

### Structural variant analysis in key salinity responsive genes

For a subset of nine genes known to be involved in salinity stress tolerance (**15, 16**), we analyzed the presence of any type of SVs within any of the paralog copies of the gene or in their putative promoter regions (i.e., up to 3,000 bp upstream of their 5’ ends), (Table 4). Altogether, 16 SVs were identified (seven insertions and nine deletions), and the paralogs of seven gene families have at least one SV potentially affecting gene transcription. The identified SVs has a length ranging from 56 bp to 4,740 bp. Eight of them are associate to DNA TEs, one is an LTR-RT, two are uncharacterized TEs, one is a microsatellite and the remining four are uncharacterized. Nine of these SVs are located inside the hosting gene (in intron). Two of the genes didn’t show any insertional polymorphism related to SVs.

**Table 4.**
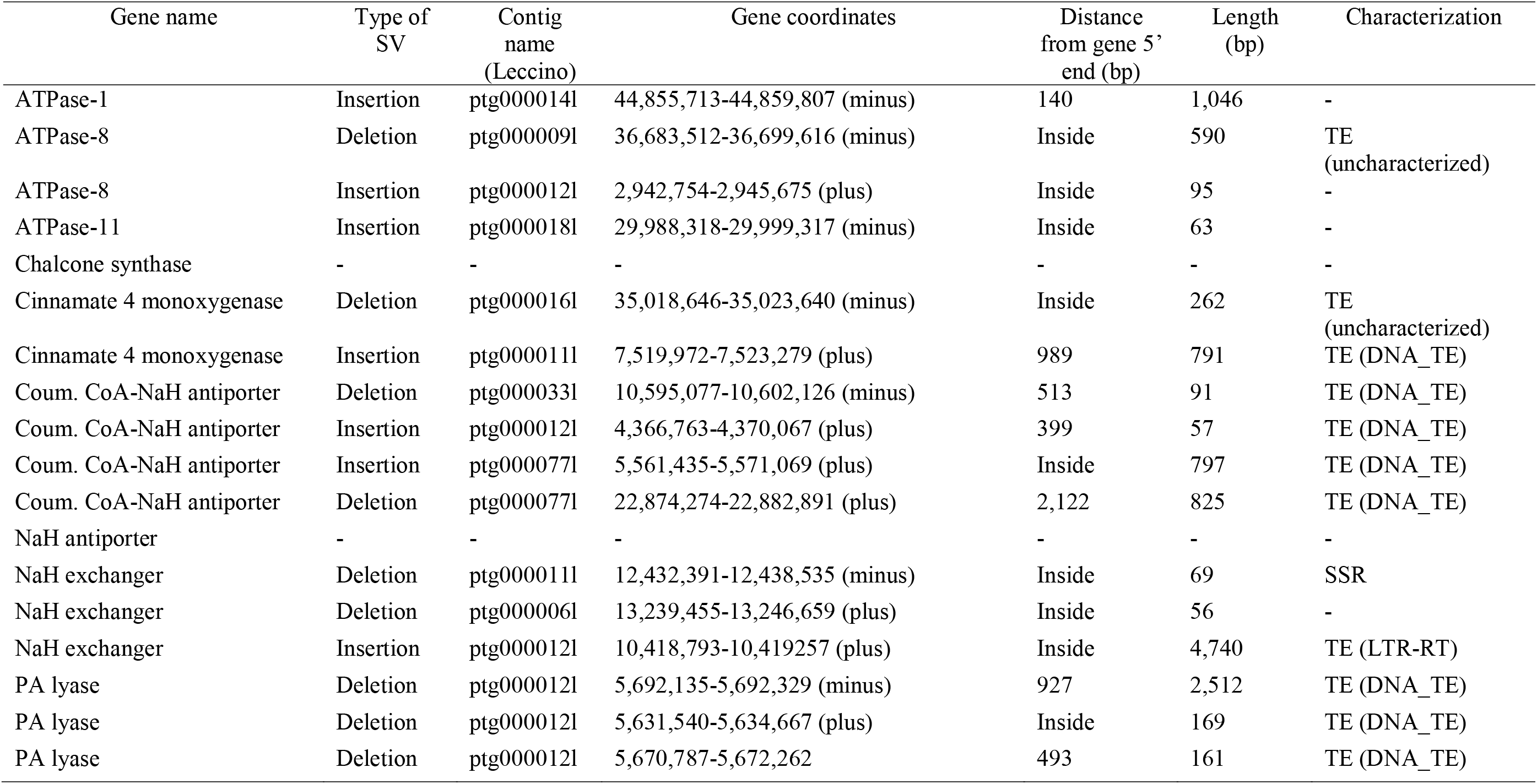
Structural variant analysis in salt related genes of ‘Frantoio’ and ‘Leccino’. Leccino is used as reference genome. Coum. = Coumarate.

The data on SVs found for *ATPase11* and *ATPase8* genes in ‘Frantoio’, could suggest a causal mechnism to explain the finding of Sodini *et al*. (**15**), i.e. an overexpression of these genes only in the salt tolerant cultivar ‘Frantoio’ when both cultivars were treated with 120 mM NaCl. Possibly the SVs associate with these genes could have a role in ‘Frantoio’ tolerance to salinity, a hypothesis that deserve further investigation. Regarding the Na+/H+ antiporter (*SOS1*) the lack of SVs between ‘Frantoio’ and ‘Leccino’ rule out their potential involvement in the overexpression of these genes in ‘Frantoio’ under salinity.

In conclusion, the assembly of ‘Frantoio’ and ‘Leccino’ provided in this study could be a valuable dataset for studying cultivar differentiation in traits (i.e. salinity) that are useful for olive genetic improvement. Together with others published genomes will enable further understanding of olive evolution and domestication processes.

## Data Records

All the raw sequencing data and the genome assemblies hve been submitted to NCBI under the bioproject accession numbers PRJNA1197703 and PRJNA1197712.

A copy of the present manuscript has been deposited at BioRxiv (https://www.biorxiv.org/) preprint archive.

## Technical Validation

The quality and concentration of extracted DNA were assessed using NanoDrop Spectrophotometer and FEMTO size profile before the genome sequencing.

After the genome assembly was completed, the assembly results were evaluated:

i. The HiFi reads used for genome assemblies were mapped back onto to the assembled genome. The alignment rate of reads was in both cases higher than 99%, showing high consistency between the reads and assembled genomes.
ii. the completeness of the genome assemblies was evaluated using the BUSCOs eudicotyledons_odb10 database.

## Code availability

Software and pipelines were executed according to the manual and protocols.

## Acknowledgements

This study was carried out within the Agritech National Research Center and received funding from the European Union Next-GenerationEU (PIANO NAZIONALE DI RIPRESA E RESILIENZA (PNRR)–MISSIONE 4 COMPONENTE 2, INVESTIMENTO 14–DD 1032 17/06/2022, CN00000022). This manuscript reflects only the authors’ views and opinions, neither the European Union nor the European Commission can be considered responsible for them. Iqra Sarfraz Agrobioscience PhD Scholarship was funded by Programma Operativo Nazionale Ricerca e Innovazione 2014-2020 (CCI 2014IT16M2OP005), FSE REACT-EU, Azione IV.4 “Dottorati e contratti di ricerca su tematiche dell’innovazione” e Azione IV.5 “Dottorati su tematiche Green”.

## Author information

### Authors contributions

L.S: Conceptualization, Methodology, Resources, Formal analysis, Data curation, Software, Writing-original draft, Project administration, Funding acquisition, Writing-review & editing.

I.S.: Methodology, Investigation, Validation, Formal analysis, Data curation, Software, Visualization, Writing-original draft, Writing-review & editing.

A.Z.: Conceptualization, Methodology, Validation, Resources, Formal analysis, Data curation, Software, Visualization, Funding acquisition, Writing-review & editing.

A.F.: Conceptualization, Writing-review & editing

M.C.: Methodology, Validation, Data curation, Software, Visualization, Writing-review & editing.

R.A.W: Methodology, Resources, Writing-review & editing

## Ethics declarations and competing interests

The authors declare no competing interests.

