## Supplementary for "PacBio genome assembly of *Olea europaea* L. subsp. *europaea* cultivars ‘Frantoio’ and ‘Leccino’ reveal main structural differences in key genes related to salt stress"

^2^[King Abdullah University of Science & Technology](https://scholar.google.com/citations?view_op=view_org&hl=en&org=15282493088823841469) (KAUST), Saudi Arabia

Rod A. Wing

**Contents**

| **Title** | **Page** |
| --- | --- |
| Supplementary Figure 1. | 2 |
| Supplementary Figure 2. | 3 |
| Supplementary Figure 3. | 4 |
| Supplementary Figure 4. | 5 |
| Supplementary Figure 5. | 6 |
| Supplementary Figure 6. | 7 |
| Supplementary Figure 7. | 8 |
| Supplementary Figure 8. | 9 |
| Supplementary Table 1. | 10 |
| Supplementary Table 2. | 11 |

**A)**
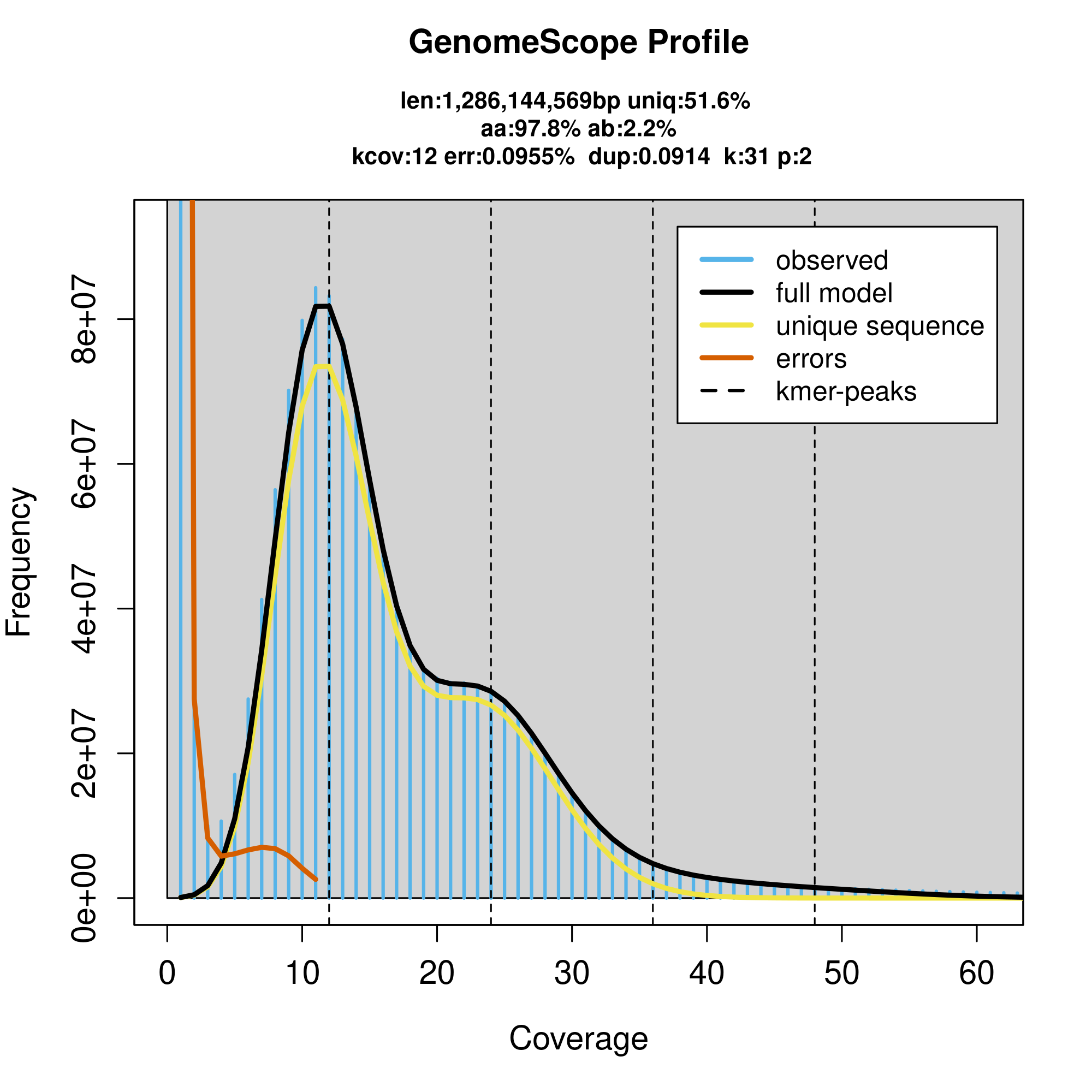
**B)**
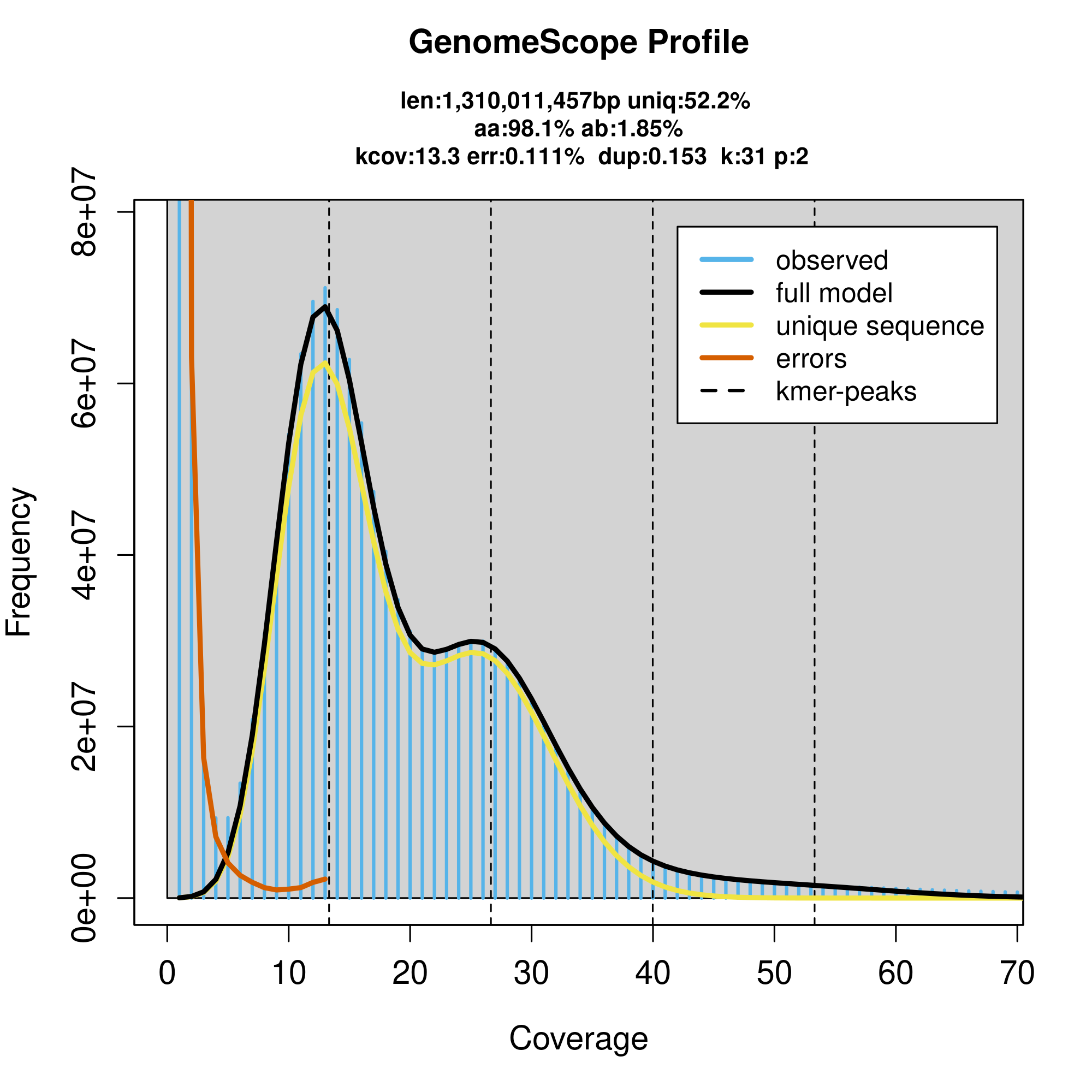

**Supplementary Figure 1**. Olea europaea L. subsp. europaea cv ‘Frantoio’ (A) and ‘Leccino’ (B) genome size estimation (len) using Jellyfish and GenomeScope2 k-mer (31) displaying homozygosity (aa), heterozygosity (ab), mean k-mer coverage for heterozygous bases (kcov), read error rate (err), the average rate of read duplications (dup), k-mer size used on the run (k), and ploidy (p).

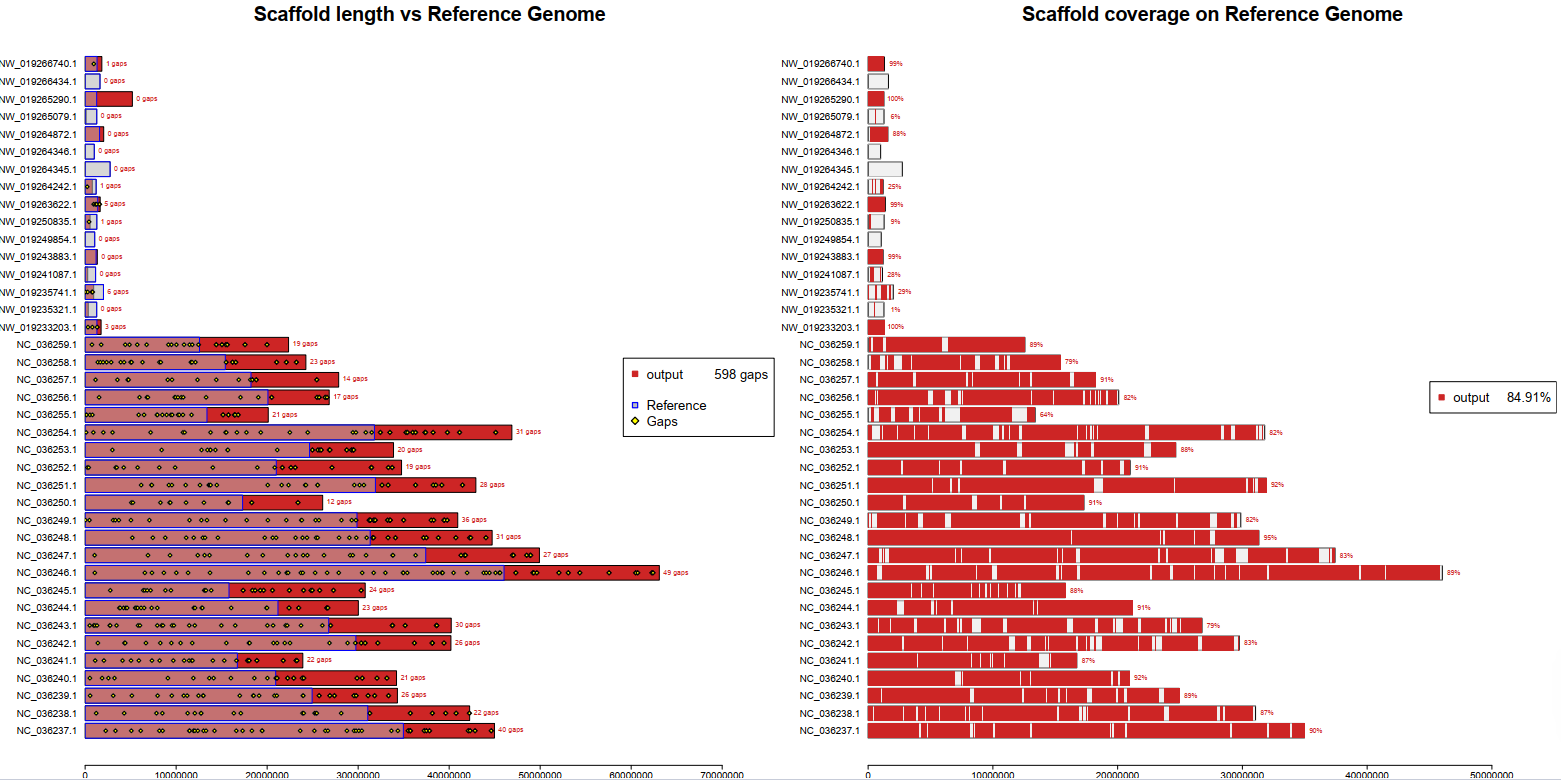

**Supplementary Figure 2: ‘**Frantoio’ scaffolding using ‘Sylvestris’ (*Olea europaea* subsp. *sylvestris*) as a reference genome.

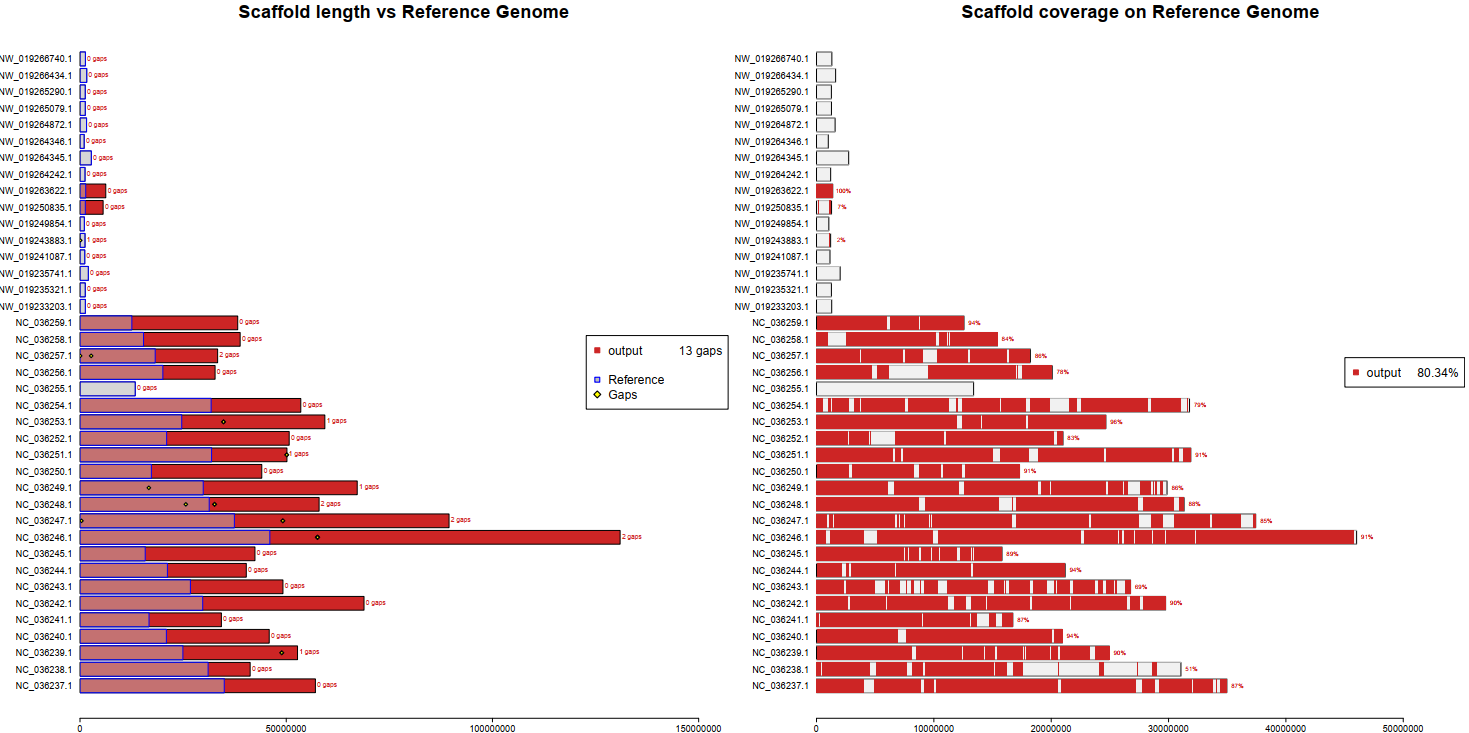

**Supplementary Figure 3: ‘**Leccino’ scaffolding using ‘Sylvestris’ (*Olea europaea* subsp. *sylvestris*) as a reference genome.

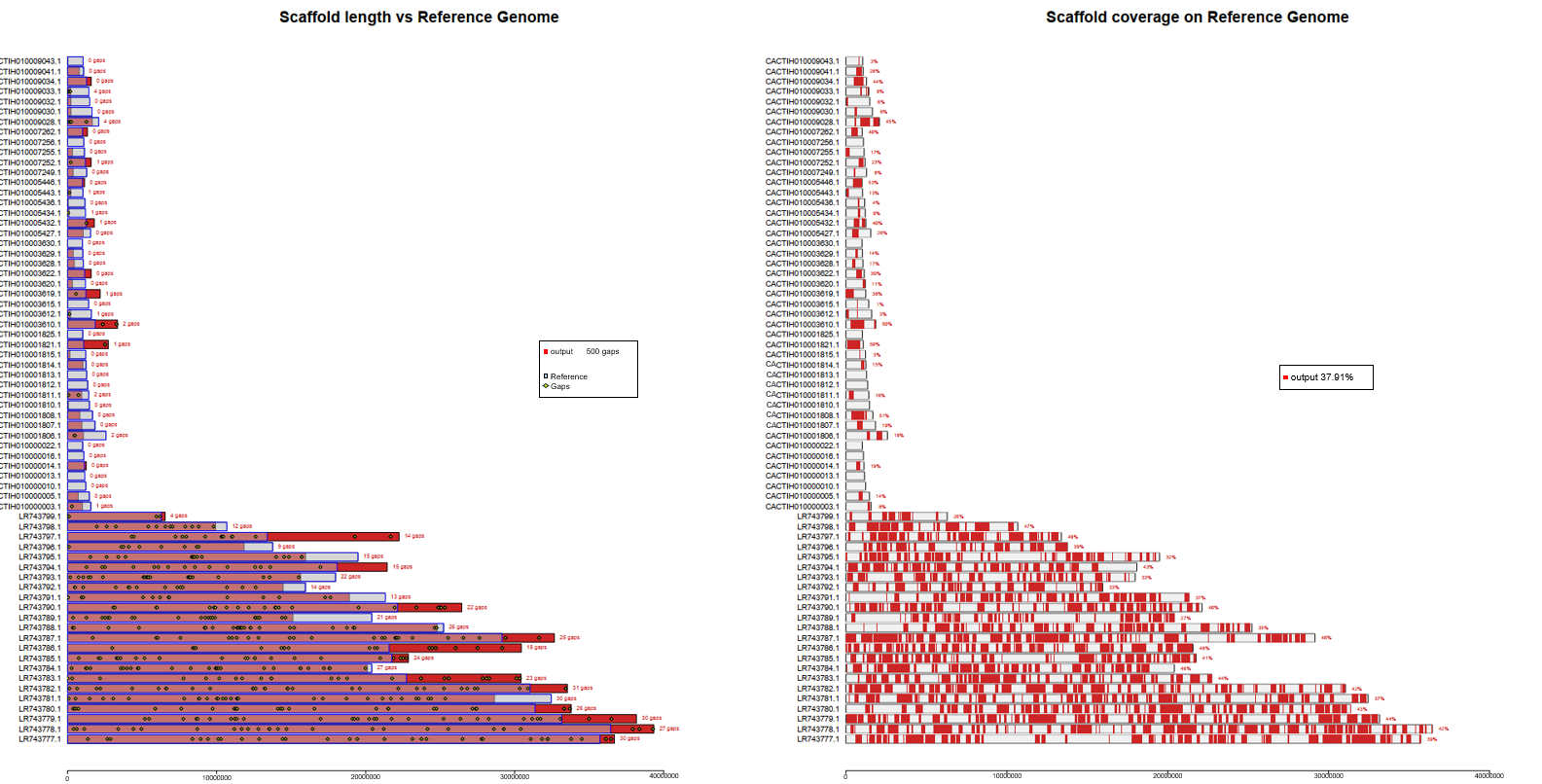

**Supplementary Figure 4: ‘**Frantoio’ scaffolding using ‘Farga’ (*Olea europaea* subsp. *europaea*) as a reference genome

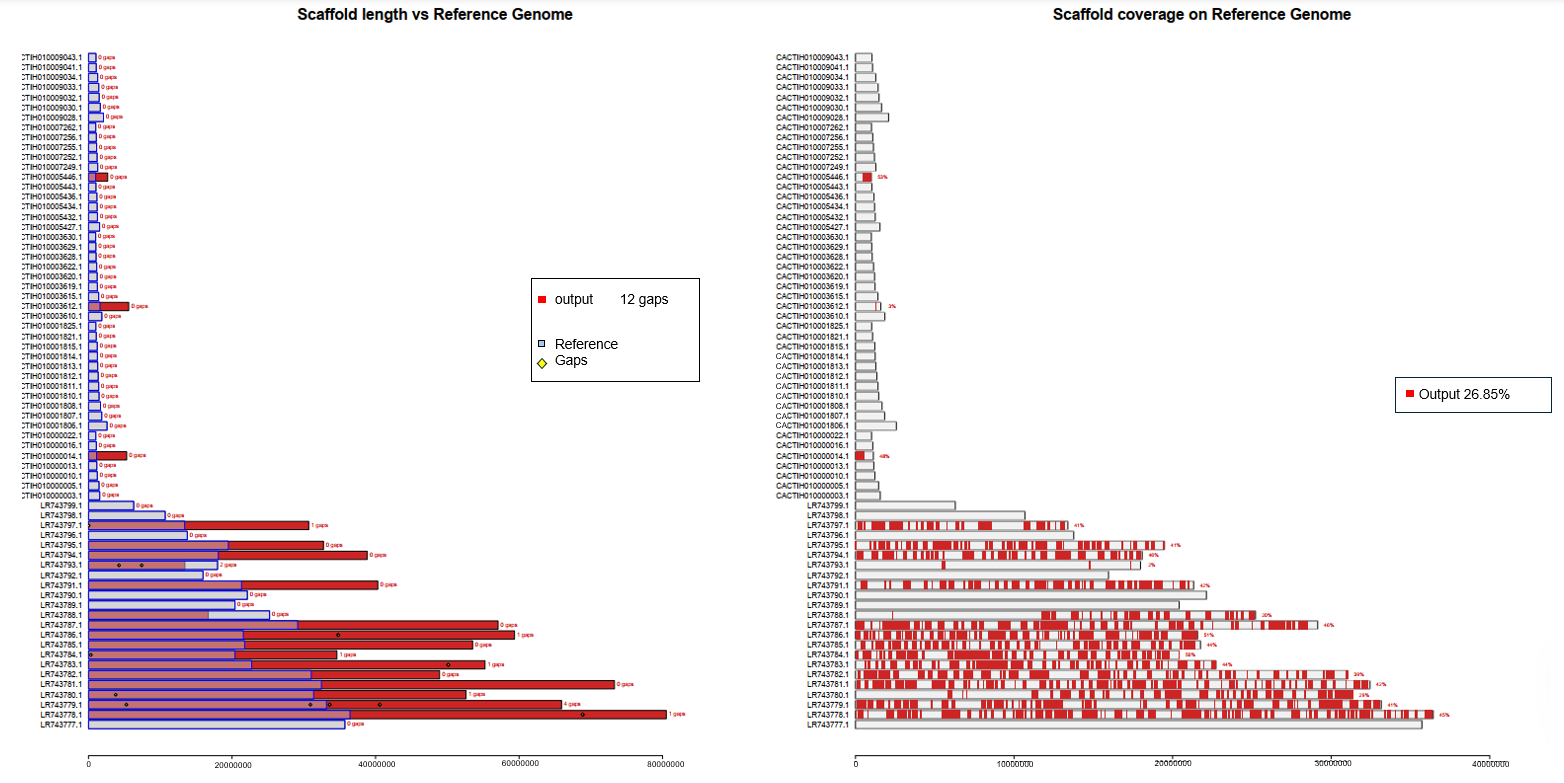

**Supplementary Figure 5: ‘**Leccino’ scaffolding using ‘Farga’ (*Olea europaea* subsp. *europaea*) as a reference genome.

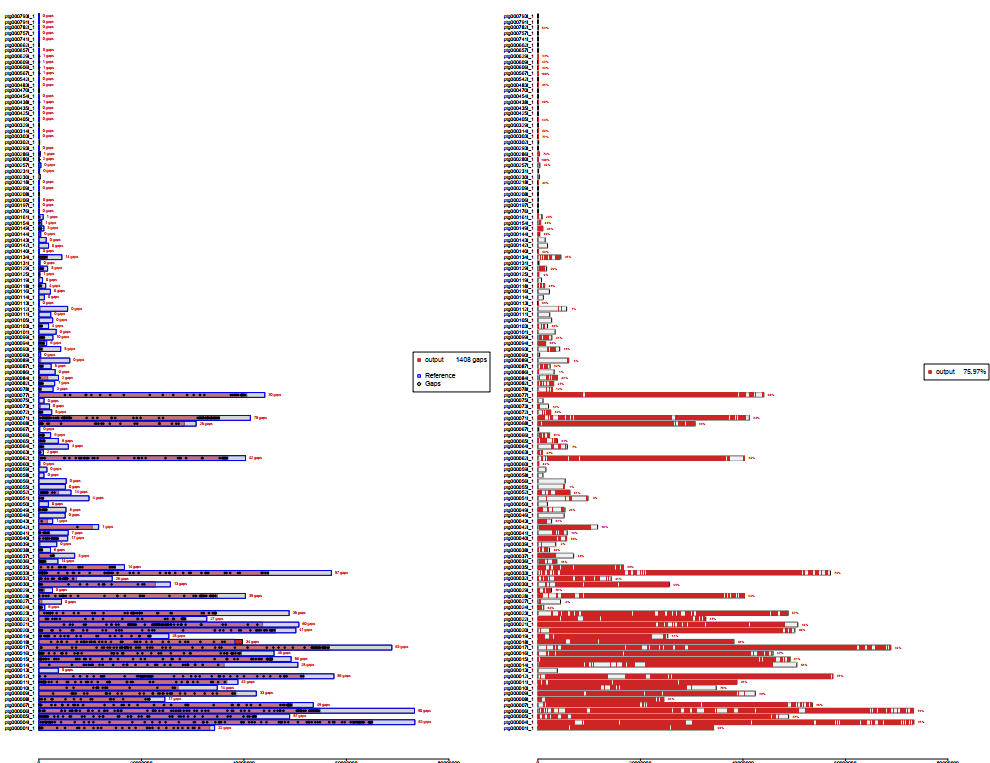

**Supplementary Figure 6: ‘**Frantoio’ scaffolding using ‘Leccino’ (this study data) as a reference genome

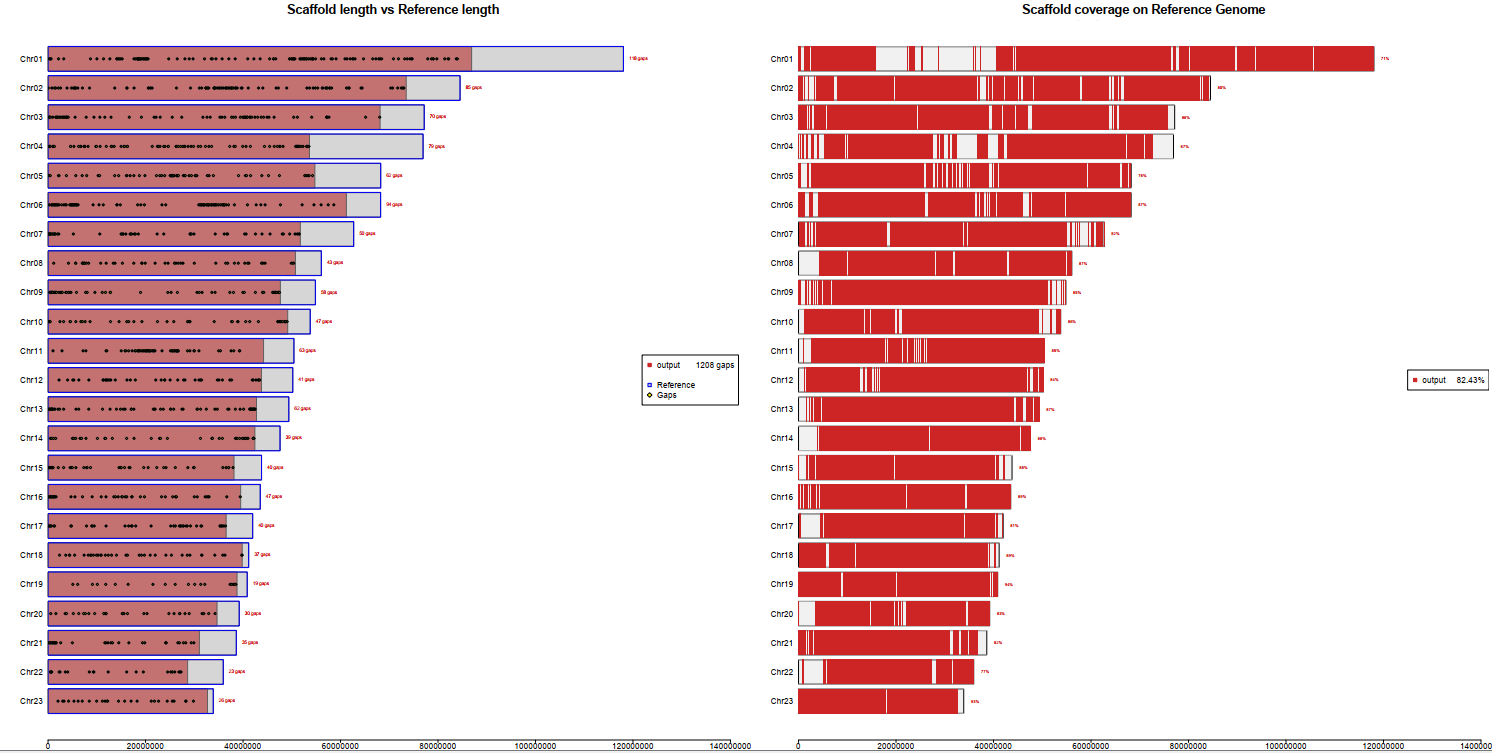

**Supplementary Figure 7: ‘**Frantoio’ scaffolding using ‘Leccino’ (Lv et al., 2024 data) as a reference genome.

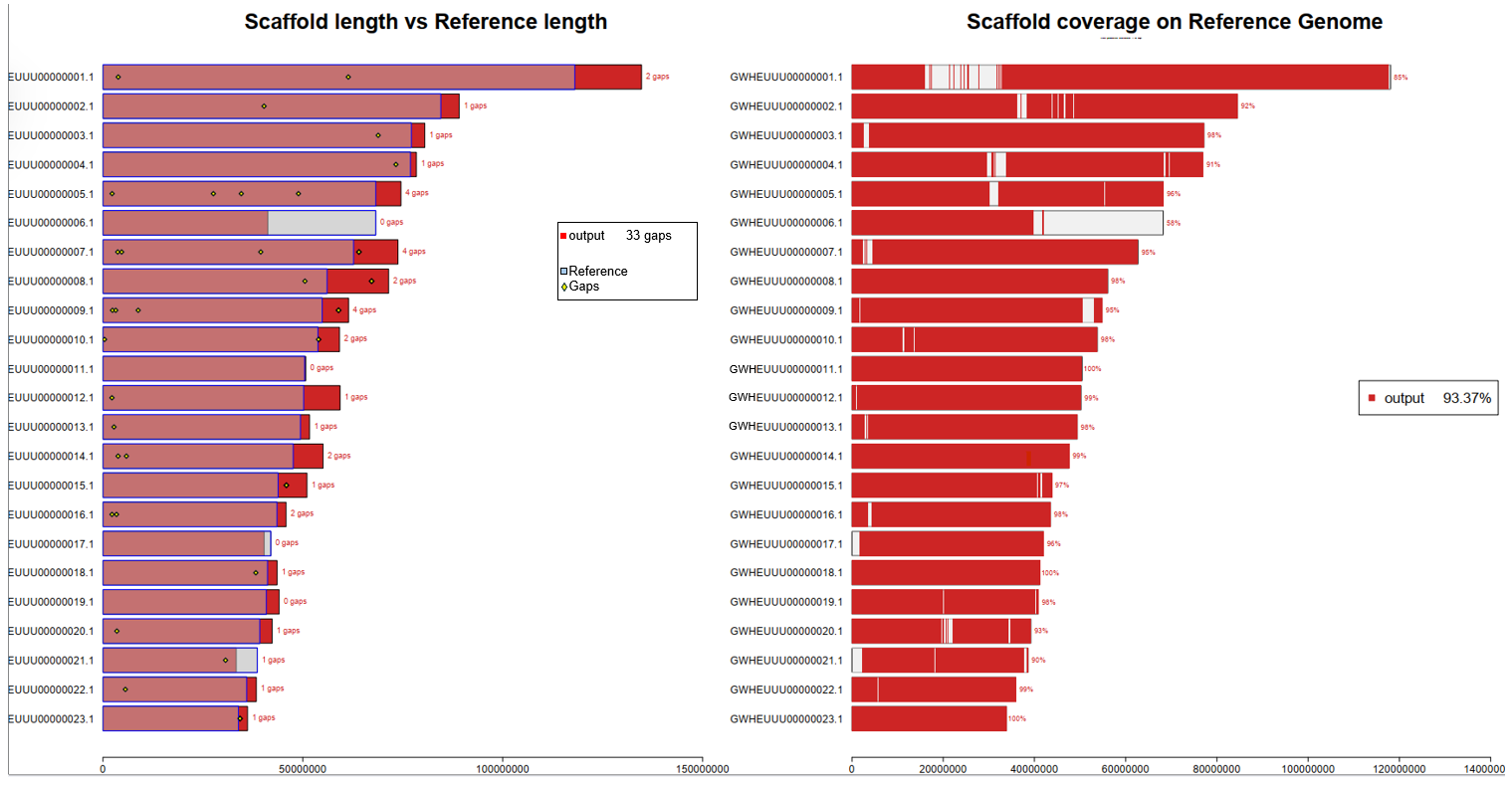

**Supplementary Figure 8:** ‘Leccino’ assembly (this study data) scaffolding using ‘Leccino’ (Lv et al., 2024 data) as a reference genome.

**Supplementary Table 1**. BUSCO main statistics as percentage (%) of the total (n=1,614) BUSCO groups searched in *Olea europaea* L. genome cultivars ‘Frantoio’ and ‘Leccino’.

| Parameter | Frantoio | Leccino |
| --- | --- | --- |
| Single copy (%) | 83.1 | 82.9 |
| Duplicated (%) | 14.7 | 15.1 |
| Fragmented (%) | 1.4 | 1.4 |
| Missing (%) | 0.8 | 0.6 |

**Supplementary Table 2.** The sequences of the three most abundant satellite repeats identified in ‘Frantoio’ and ‘Leccino’ genomes.

>Satellite_1

GTTTCCAATCAACCTGCCAAATTTCACCCCATTCTGACACCGTCGCGAAAAATGTCGAAA

TTGCCCCTAGCGCGTTTTTT

>Satellite_2

GTGCCGGTTCTTCCACATGAACCTCGTTCTTAATCGTCATGAGCTTTTCGAAATGTGGTT

ATGGTACCAAAACGGACATAGTATGCAAAAGTTACGCCAATTTTTTCGAAAACTAACTAT

TTAAGCATTCCGTGTACATAC

>Satellite_3

ATTCCAATCGAAATAGACTGCCAATTCGCCAAAATGATCAAACTCGAAGATATCATGGAA

TCAAAAGCCACTTTTCATTGGCTCGATTCCATTCGAATTAGACGAGC
